# Zoonotic tuberculosis in India: looking beyond *Mycobacterium bovis*

**DOI:** 10.1101/847715

**Authors:** Shannon C Duffy, Sreenidhi Srinivasan, Megan A Schilling, Tod Stuber, Sarah N Danchuk, Joy S Michael, Manigandan Venkatesan, Nitish Bansal, Sushila Maan, Naresh Jindal, Deepika Chaudhary, Premanshu Dandapat, Robab Katani, Shubhada Chothe, Maroudam Veerasami, Suelee Robbe-Austerman, Nicholas Juleff, Vivek Kapur, Marcel A Behr

## Abstract

**Background:** Zoonotic tuberculosis (zTB) is the transmission of *Mycobacterium tuberculosis* complex (MTBC) subspecies from animals to humans. zTB is generally quantified by determining the proportion of human isolates that are *Mycobacterium bovis*. Although India has the world’s largest number of human TB cases and the largest cattle population, where bovine TB is endemic, the burden of zTB is unknown.

**Methods:** To obtain estimates of zTB in India, a PCR-based approach was applied to sub-speciate positive MGIT® cultures from 940 patients (548 pulmonary, 392 extrapulmonary disease) at a large referral hospital in India. Twenty-five isolates of interest were subject to whole genome sequencing (WGS) and compared with 715 publicly available MTBC sequences from South Asia.

**Findings:** A conclusive identification was obtained for 939 samples; wildtype *M. bovis* was not identified (95% CI: 0 – 0.4%). There were 912 *M. tuberculosis sensu stricto* (97.0%, 95% CI: 95.7 – 98.0), 7 *M. orygis* (95% CI: 0.3 – 1.5%); 5 *M. bovis* BCG, and 15 non-tuberculous mycobacteria. WGS analysis of 715 MTBC sequences again identified no *M. bovis* (95% CI: 0 – 0.4%). Human and cattle MTBC isolates were interspersed within the *M. orgyis* and *M. tuberculosis sensu stricto* lineages.

**Interpretation:** *M. bovis* prevalence in humans is an inadequate proxy of zTB in India. The recovery of *M. orygis* from humans, together with the finding of *M. tuberculosis* in cattle, underscores the need for One Health investigations to assess the burden of zTB in countries with endemic bovine TB.

**Funding:** Bill & Melinda Gates Foundation, Canadian Institutes for Health Research

## Introduction

Tuberculosis (TB) is the world’s most deadly disease by a single infectious agent, claiming over 1.4 million lives each year, primarily in low-and middle-income countries (LMICs) (^1^). Tuberculosis in cattle (bovine TB or bTB) is also a significant animal health problem endemic in most LMICs, costing an estimated 3 billion USD worldwide each year (^2^). The World Health Organization (WHO), the World Organisation for Animal Health (OIE), and the Food and Agriculture Organization for the United Nations (FAO) define zoonotic TB (zTB) as human infection with *Mycobacterium bovis*, a member of the *Mycobacterium tuberculosis* complex (MTBC) (^1,3,4^). Therefore, the detection of *M. bovis* is most often used as a proxy for measuring zTB prevalence. Based on this definition, the WHO has recently estimated that the annual number of human TB cases due to zTB is 143,000 (^1^).

The WHO aims to reduce TB incidence by 90% by 2035 as a part of the End TB Strategy (^5^). India carries the world’s largest burden of human TB, with over 2.6 million cases and 400,000 deaths reported in 2019 (^1^). The cattle population in India exceeds 300 million, a population larger than any other country (^6^). Yet, bovine TB is both uncontrolled and endemic, with an estimated 21.8 million infected cows in India (^7^). A small number of previous studies estimated the prevalence of zTB in India to be ~10%, however these studies were limited in their abilities to differentiate between MTBC members (^8–10^). Recent evidence has suggested that *M. orygis*, a newly described MTBC member, may be endemic to South Asia (^11–14^). However, robust estimates of *M. orygis* prevalence in humans or cattle are lacking.

In this study, we aimed to begin to obtain an estimate of the prevalence of zTB in India through molecular analyses of 940 positive broth cultures from a large referral hospital in Southern India. To evaluate our findings within the framework of other sequences collected from South Asia, we subjected twenty-five isolates to whole genome sequencing (WGS) and compared these sequences to representative genomes of the MTBC lineages circulating in the world as well as 715 publicly available genomes collected from cattle and humans from South Asia.

## Methods

### Study design

Positive MGIT® cultures collected at Christian Medical College (CMC) in Vellore, Tamil Nadu were screened using PCR assays previously established and validated at McGill University, Montreal, Quebec. Institutional Review Board (IRB Min. No. 11725 dated December 19^th^, 2018) and ethical clearance were obtained to perform this study. All patient data was denominalized prior to inclusion of each isolate. This study took place from January 2019 to June 2019. WGS was performed at Lala Lajpat Rai University of Veterinary and Animal Sciences (LUVAS) in Hisar, Haryana. WGS analysis and construction of phylogenetic trees was performed at The Pennsylvania State University, State College, Pennsylvania, in collaboration with the United States Department of Agriculture in Ames, Iowa. A flowchart of study design is provided in supplementary figure 1.

### Sample collection

MGIT® cultures were obtained from patients from 20 states and territories in India as well as Bangladesh and Nepal who visited the outpatient department at CMC with signs and symptoms of probable tuberculosis from October 2018 – March 2019. In total, 548 pulmonary and 392 extrapulmonary MGIT ® cultures were collected for screening in this study. Pulmonary isolates were defined as those collected from the lung fluid or tissue. This included sputum, lung biopsies, bronchoalveolar lavage, endotracheal aspirate, pleural samples, and one chest wall abscess. Extrapulmonary isolates were defined as those collected from tissue other than the lungs. The extrapulmonary samples types are listed in supplementary table 1. This definition serves to differentiate disseminated disease from disease confined to the lungs and their adjacent structures.

### Conventional PCR

All PCR screening was performed at CMC, Vellore using DNA prepared by boiling. The first 600 isolates were screened in January 2019 by conventional PCR to assess the proportion of *M. tuberculosis sensu stricto*, as opposed to *M. bovis*, *M. orygis* or other MTBC subspecies. Two deletion-based conventional PCR assays were developed to screen the first 600 samples for zTB. A three-primer PCR was designed to detect the presence or absence of region of difference 9 (RD9), which is present in *M. tuberculosis* and absent from other MTBC subspecies (Supplementary Figure 2A, B). A six-primer PCR was developed to detect differences in the deletion size of RD12. The RD12 region is present in *M. tuberculosis* and absent from *M. bovis* and *M. orygis.* However, the *M. orygis* deletion is larger and IS*6110* is inserted (Supplementary Figure 2C, D). All primers were designed using the Primer3 software v. 0.4.0 (Supplementary Table 2) (^15^). PCR reactions were performed as described in supplementary table 3. PCR products were separated by gel electrophoresis on a 2% (wt/vol) agarose gel containing a 1:10K dilution of ethidium bromide.

### Real-time PCR

A five-probe multiplex real-time PCR assay was developed to screen the remaining 340 isolates in May 2019. Forty isolates identified as *M. tuberculosis* and all isolates previously identified as not *M. tuberculosis sensu stricto* by conventional PCR were also assessed by real-time PCR to confirm consistent identity across assays. Four probes were previously described by Halse *et al* (^16^), which detect the presence of RD1, RD9, RD12, and ext-RD9. A fifth probe was designed to target the *M. orygis*-specific single nucleotide polymorphism (SNP) in *Rv0444c* g698c (^17^). These probes allowed for differentiation of several members of the MTBC (Supplementary Table 4). The Rv0444c probe and primers were designed using Primer Express v. 3.0 (Applied Biosystems). All primer and probe sequences are listed in supplementary table 2. Real-time PCR reactions were performed as described in supplementary table 3.

### Whole genome sequencing

Isolates that were identified as MTBC other than *M. tuberculosis sensu stricto* or as inconclusive by PCR were processed for WGS. Extraction of genomic DNA was performed as previously described (^18^). Libraries were prepared using the Nextera DNA flex library prep kit (Illumina). The quality of each library was assessed using a Fragment Analyzer™ automated CE system (Agilent) using an NGS fragment kit (1-6000bp). Libraries were paired-end sequenced on an Illumina MiSeq using the MiSeq Reagent Kit v3, 600-cycle.

### hsp65 *sequencing*

For isolates where no amplification occurred on PCR screening, suggestive of a non-tuberculous mycobacteria (NTM), *hsp65* sequencing was done by the Sanger method, as previously described (^19^).

### Bioinformatics

Genome sequences were assessed using the United States Department of Agriculture Animal and Plant Health Inspection Service Veterinary Services pipeline vSNP (https://github.com/USDA-VS/vSNP). The vSNP pipeline involved a two-step process. Step 1 determined SNP positions called within the sequence. Step 2 assessed SNPs called between closely related isolate groups to output SNP alignments, tables and phylogenetic trees. For a SNP to be considered in a group there must have been at least one position with an allele count (AC) =2, quality score >150 and map quality > 56. The output SNP alignment was used to assemble a maximum likelihood phylogenetic tree using RAxML (^20^). Additional details of the pipeline are provided in supplementary methods.

### Phylogenetic tree assembly

To compare the newly sequenced genomes to sequences from South Asia, a NCBI Sequence Read Archive (SRA) (https://www.ncbi.nlm.nih.gov/sra) search was performed using the search terms (“*Mycobacterium bovis*” OR “*Mycobacterium tuberculosis*” OR “*Mycobacterium africanum*” OR “*Mycobacterium orygis*” OR “*Mycobacterium canetti*” OR “*Mycobacterium caprae*” OR “*Mycobacterium bovis* BCG” NOT “H37Rv” NOT “H37Ra”) from (“India” OR “Bangladesh” OR “Nepal” OR “Sri Lanka” OR “Pakistan”). All sequences were downloaded from the SRA using the fasterq-dump tool from the sra toolkit v. 2.9.6 (https://ncbi.github.io/sra-tools/) and sequences were then filtered for quality, according to the criteria detailed in supplementary figure 3. Sequences that met the indicated quality were run through vSNP (Supplementary Table 5). Phylogenetic trees were constructed using vSNP to compare the sequences from this study with the genomes from South Asia. Reference sequences were also included for comparison (Supplementary Table 6). Phylogenetic trees were rooted to *M. tuberculosis* H37Rv. To compare the sequences collected in this study in the context of the global MTBC, treeSPAdes (http://cab.spbu.ru/software/spades/) was used to assemble reads for kSNP3 (^21^). The kSNP3 manual instructions were followed using kchooser calculated kmer value. All phylogenetic trees were visualized using the Interactive Tree of Life (iTOL) with their respective metadata (Supplementary Table 7) (^22^). Additional description of tree assembly is provided in supplementary methods.

### Statistics

Statistical analysis was performed using GraphPad Prism 8.1.2. Confidence intervals were calculated using the binomial exact calculation. Group comparisons were assessed using a Fisher’s exact test where p < 0.05 was considered significant. Relative risk (RR) confidence intervals values were calculated using a Koopman asymptotic score.

## Results

### Sample characteristics

DNA was prepared from a total of 940 positive MGIT® cultures. In total, 548 pulmonary and 392 extrapulmonary isolates were included in this study. Table 1 describes the characteristics of the patients from whom the isolates were obtained. Extrapulmonary TB occurred more often in younger ages. In patients <10 years old, the relative risk (RR) of extrapulmonary TB was 1.8 (95% CI: 1.3 – 2.2, p = 0.002). Pulmonary TB occurred more often in older ages. In patients of ≥60 years, the relative risk of pulmonary TB was 1.4 (95% CI = 1.3 – 1.6, p < 0.0001). The patients were from 20 states and territories in India (N=884), from Bangladesh (N=54), and from Nepal (N=2) (Figure 1A). Within India, the majority of isolates were collected from patients living in Tamil Nadu, (N=341), West Bengal (N=187), and Andhra Pradesh (N=100). The majority of isolates collected from each location were pulmonary in origin, with the exception of Bangladesh where 64.8% of samples were extrapulmonary (Figure 1B).

**Table 1:**
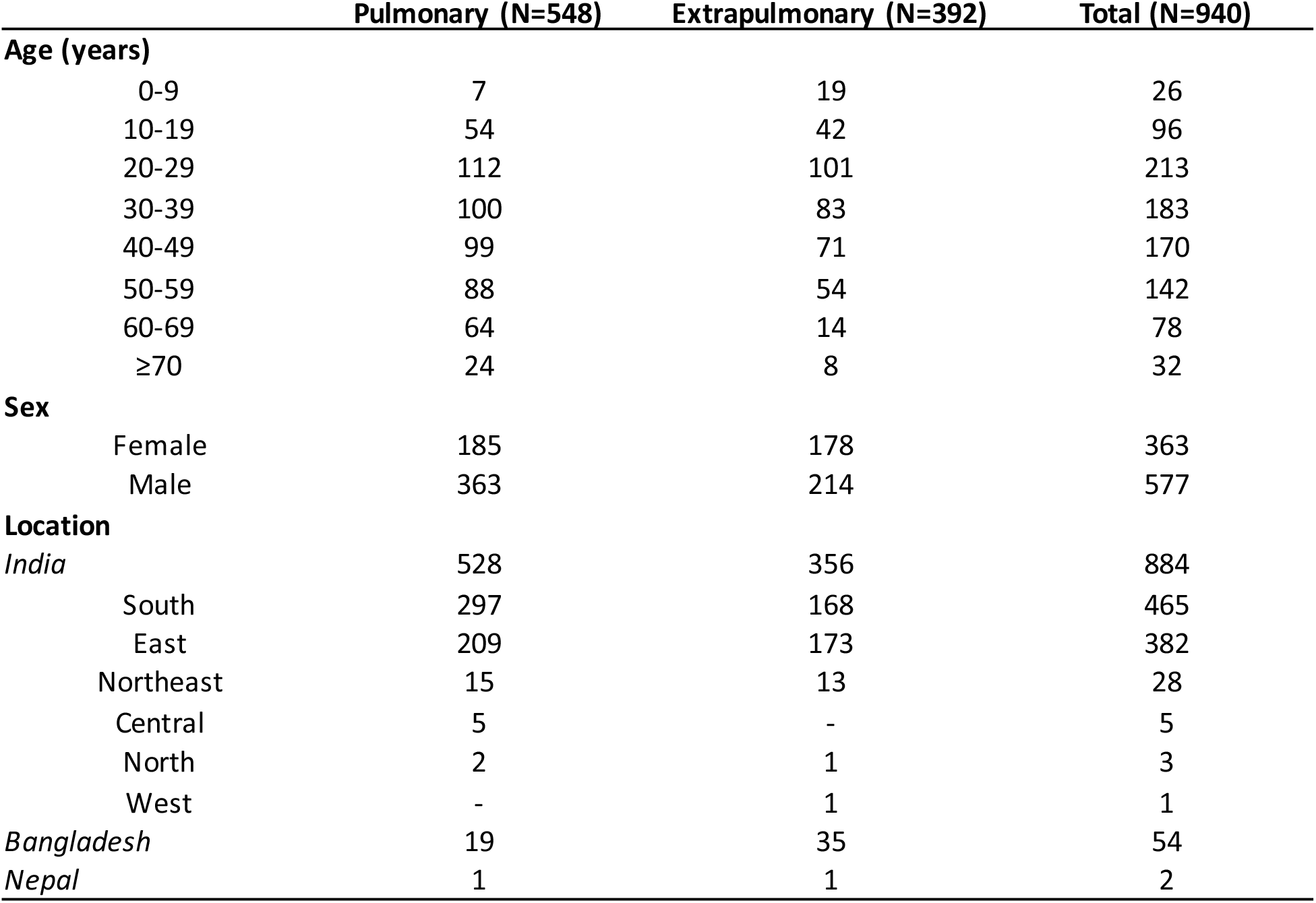
Characteristics of patients whose isolates were screened for zoonotic tuberculosis. Regions of India are divided by location in the following manner. South India includes Andaman and Nicobar Islands, Andhra Pradesh, Karnataka, Kerala, Lakshadweep, Puducherry, Tamil Nadu and Telangana. East India includes West Bengal, Bihar, Jharkhand, and Odisha. Northeast India includes Arunachal Pradesh, Assam, Manipur, Meghalaya, Mizoram, Nagaland, Sikkim, and Tripura. Central India includes Chhattisgarh and Madhya Pradesh. North India includes Jammu and Kashmir, Himachal Pradesh, Punjab, Chandigarh, Uttarakhand, Haryana, National Capital Territory of Delhi, Rajasthan, and Uttar Pradesh. West India includes Dadra and Nagar Haveli, Daman and Diu, Goa, Gujarat, and Maharashtra.

**Figure 1:**
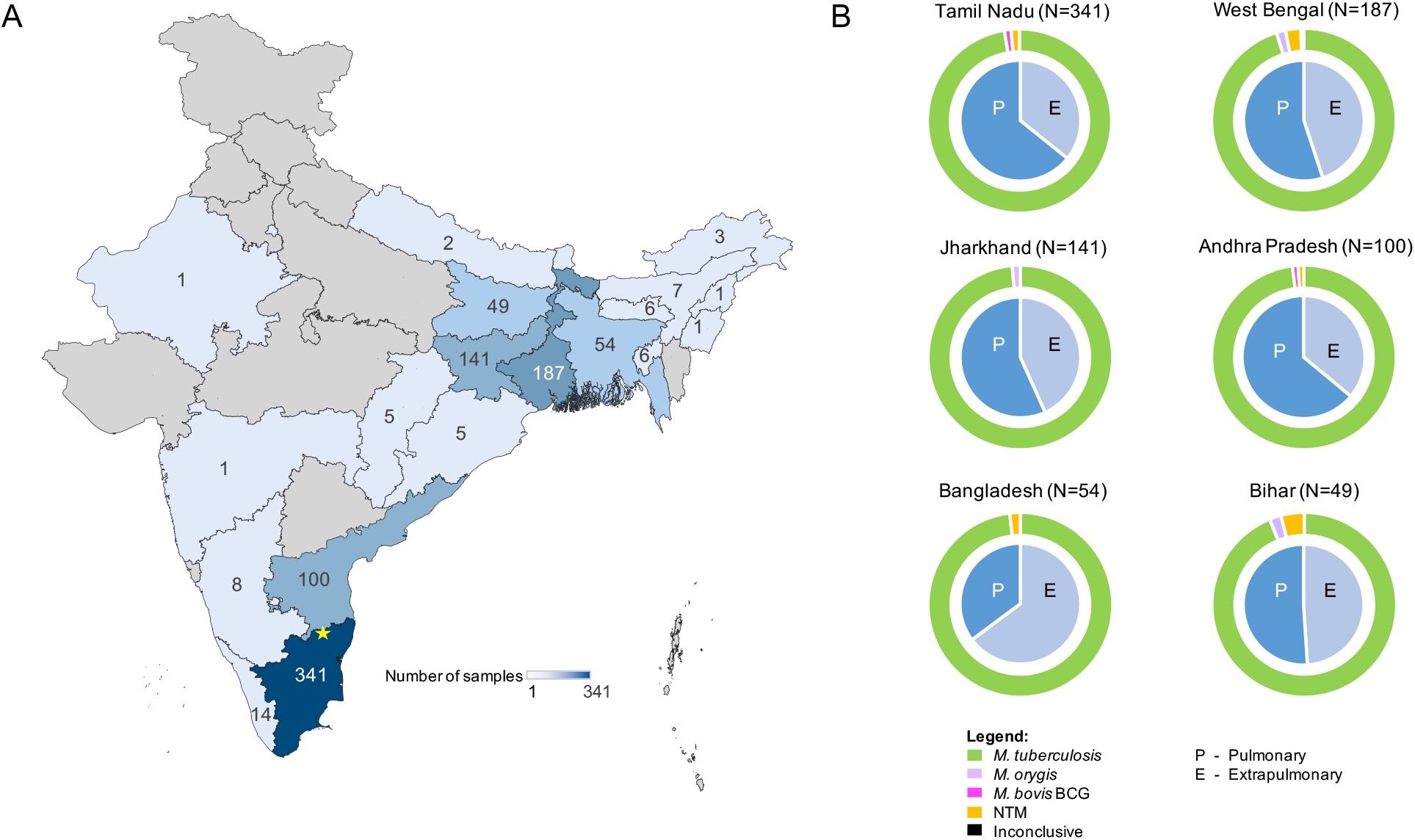
Distribution and sample types of patient isolates within India and surrounding countries Fig 1A: A representation of the geographic distribution of the collected isolates. The intensity of the color corresponds to the number of isolates collected per area. No isolates were screened from locations pictured in grey. The location of the study site (Christian Medical College, Vellore) is indicated by a yellow star. Fig 1B: Sample types of isolates from locations where 20 or more samples were collected. The inner pie chart shows the proportion of pulmonary (P) and extrapulmonary (E) isolates collected from the indicated location. The outer donut chart indicates the proportion of mycobacterial (sub)species collected from that location.

### PCR and WGS results

Isolates were assessed by conventional and real-time PCR to evaluate the proportion of MTBC subspecies. Following initial PCR screening, 25 isolates were selected for WGS (Supplementary Table 8). Based on PCR, these 25 were 7 *M. orygis*, 6 *M. bovis* BCG, 8 inconclusive, 2 *M. tuberculosis* isolates negative for RD12, and 2 additional *M. tuberculosis* isolates included erroneously. Of the 8 inconclusive isolates, 7 were so-categorized due to a delayed amplification of the RD12 probe and 1 was selected due to poor amplification of the RD1 probe.

The proportions of mycobacterial species identified following all genotyping are described in Table 2. Surprisingly, no wildtype *M. bovis* was identified in this study (0%, 95% CI: 0 – 0.4%). A total of 7 (0.7%, 95% CI: 0.3 – 1.5%) isolates were identified as *M. orygis*, 6 of which were extrapulmonary (1.5%, 95% CI: 0.6 – 3.3%). *M. orygis* was enriched in extrapulmonary isolates, where RR = 8.4 (95% CI: 1.3 – 52.9, p = 0.02). Six out of 7 *M. orygis* isolates came from patients from northeastern Indian states Bihar, Jharkhand, and West Bengal (Figure 1B). The final *M. orygis* isolate came from a patient from Karnataka. In total, 912 isolates (97.0%, 95% CI: 95.7 – 98.0%) were identified as *M. tuberculosis sensu stricto.* Two isolates lacked RD12, but WGS assigned these two as *M. tuberculosis sensu stricto* and not *M. canettii.* The 7 isolates categorized as inconclusive due to a delayed amplification of the RD12 probe contained a SNP in the RD12 reverse primer sequence. The other isolate categorized as inconclusive due to poor amplification of RD1 was found to have a 5,962bp deletion spanning Rv3871-Rv3877 which included the site of the RD1 probe. Five isolates (0.5%, 95% CI: 0.2 – 1.2%) were identified as *M. bovis* BCG. These isolates were all identified in patients ≤ 3 years old which is consistent with expectations. Fifteen isolates (1.6%, 95% CI: 0.9 – 2.6%) were identified as NTMs. The most common NTM species identified were *M. abscessus* and *M. intracellulare* (Supplementary Table 9). The identification of one isolate was inconsistent across methods suggestive of tube mislabeling: RT PCR identified it as *M. bovis* BCG and WGS identified it as *M. tuberculosis sensu stricto*.

**Table 2:**
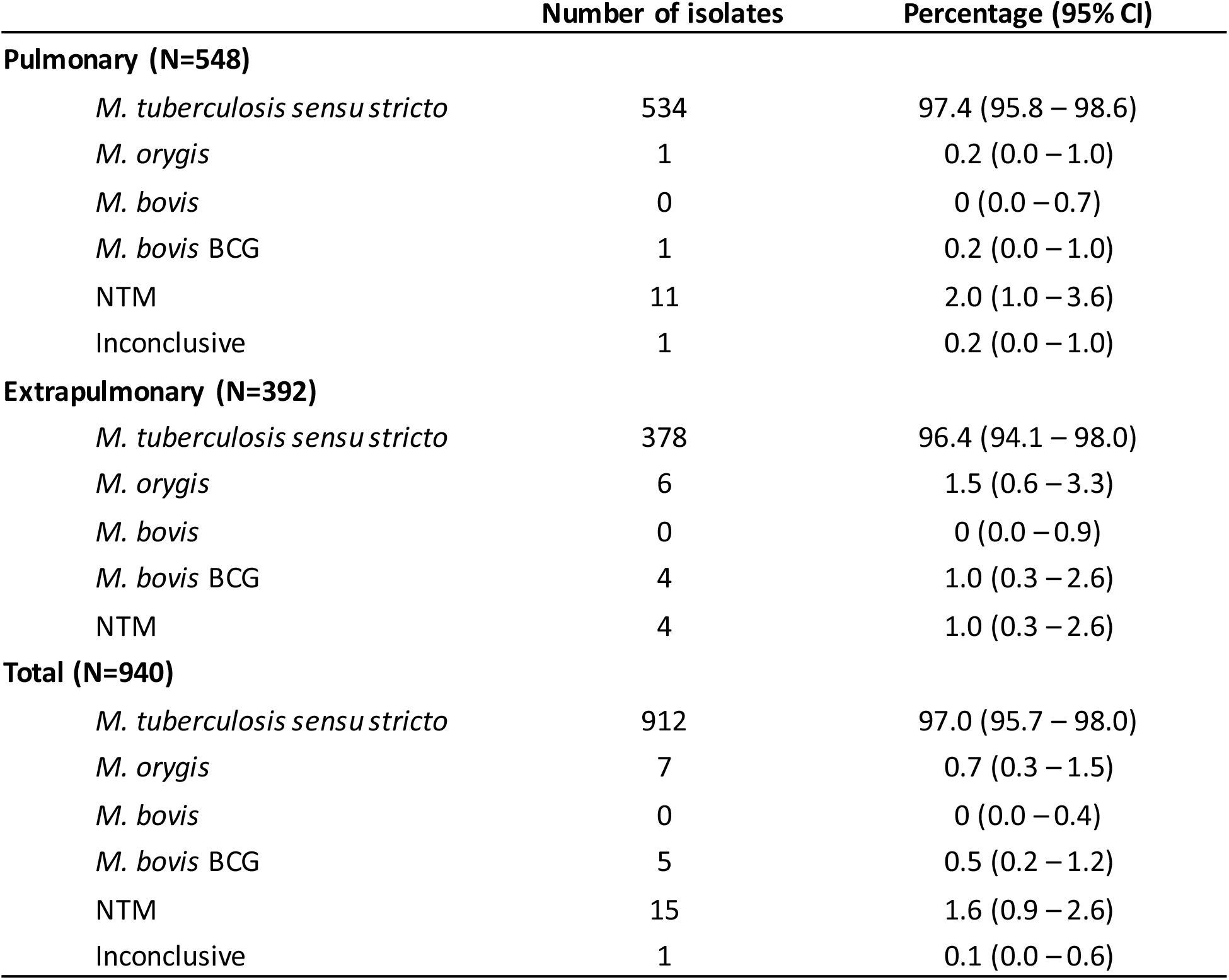
Results of PCR and WGS identification of isolates by sample type

|  | Number of isolates | Percentage (95% CI) |
| --- | --- | --- |
| **Pulmonary (N=548)** |  |  |
| *M. tuberculosis sensu stricto* | 534 | 97.4 (95.8 – 98.6) |
| *M. orygis* | 1 | 0.2 (0.0 – 1.0) |
| *M. bovis* | 0 | 0 (0.0 – 0.7) |
| *M. bovis* BCG | 1 | 0.2 (0.0 – 1.0) |
| NTM | 11 | 2.0 (1.0 – 3.6) |
| Inconclusive | 1 | 0.2 (0.0 – 1.0) |
| **Extrapulmonary (N=392)** |  |  |
| *M. tuberculosis sensu stricto* | 378 | 96.4 (94.1 – 98.0) |
| *M. orygis* | 6 | 1.5 (0.6 – 3.3) |
| *M. bovis* | 0 | 0 (0.0 – 0.9) |
| *M. bovis* BCG | 4 | 1.0 (0.3 – 2.6) |
| NTM | 4 | 1.0 (0.3 – 2.6) |
| **Total (N=940)** |  |  |
| *M. tuberculosis sensu stricto* | 912 | 97.0 (95.7 – 98.0) |
| *M. orygis* | 7 | 0.7 (0.3 – 1.5) |
| *M. bovis* | 0 | 0 (0.0 – 0.4) |
| *M. bovis* BCG | 5 | 0.5 (0.2 – 1.2) |
| NTM | 15 | 1.6 (0.9 – 2.6) |
| Inconclusive | 1 | 0.1 (0.0 – 0.6) |

### Phylogenetic analysis

A phylogenetic tree was constructed comparing the 25 isolates sequenced in this study in the context of the MTBC lineages circulating the world (Figure 2). The 13 *M. tuberculosis sensu stricto* isolates were dispersed across lineage 1 (N=9), lineage 2 (N=1), lineage 3 (N=1) and lineage 4 (N=2). Of the 9 lineage 1 genomes collected in this study, 7 were found to cluster closely together; these 7 isolates shared a SNP in the RD12 reverse primer sequence. All 5 confirmed BCG isolates identified as the Russia strain. There were no isolates that clustered with *M. bovis* or *M. caprae*. The 7 isolates that clustered with previously sequenced *M. orygis* isolates were separated by 66 – 282 SNPs from one another (Supplementary Figure 4).

**Figure 2:**
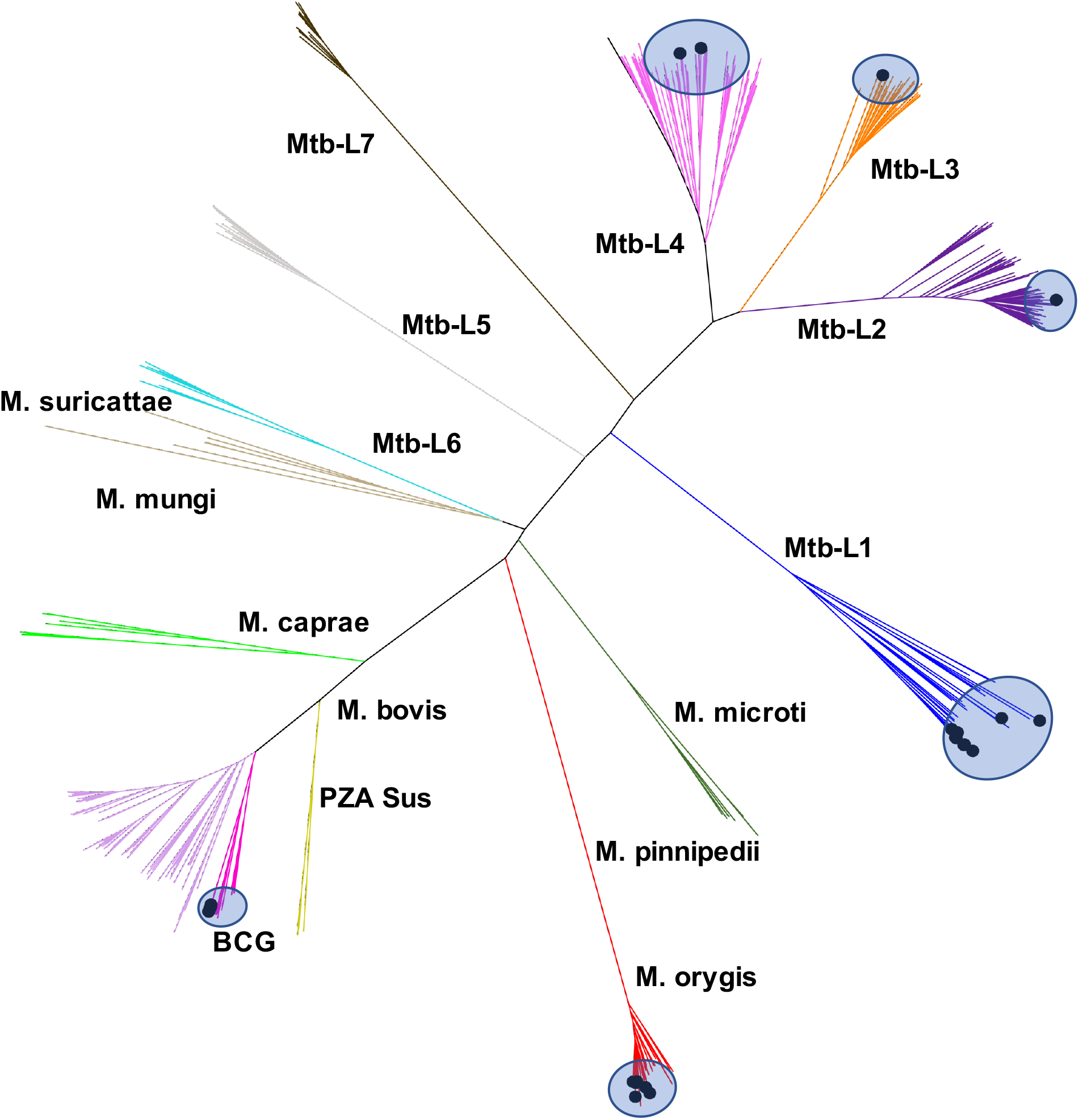
Phylogenies of newly sequenced isolates in the context of the extent of genetic diversity amongst MTBC isolates from across the world The unrooted tree shows where the isolates from this study are clustering based on representative samples from all MTBC lineages from around the world. The isolates from this study (highlighted in blue circles) are within the Mtb-L1, Mtb-L2, Mtb-L3, Mtb-L4, *M. orygis*, and *M. bovis* BCG clusters.

An SRA search for MTBC isolates from South Asia generated 1640 genomes. These sequences were then filtered prior to tree assembly to yield a total of 715 high-quality sequences (Supplementary Figure 3). The downloaded genomes were phylogenetically compared with the newly sequenced isolates and their respective metadata (Figure 3A). Again, no wild-type *M. bovis* was identified in the 715 genomes (95% CI: 0 – 0.4%). As expected, there was an enrichment of *M. tuberculosis* lineage 1 and 3. Almost all lineage 1 samples were from India whereas the lineage 2-4 sequences were collected from a variety of other countries in South Asia. Cattle origin sequences from South Asia were represented in *M. tuberculosis* lineage 1 and the *M. orygis* lineage. The phylogenetic relationships of the isolates from this study suggest that the 4 cattle origin isolates in lineage 1 were dispersed among human sequences; likewise, the 7 human *M. orygis* isolates were dispersed among sequences collected from cattle (Figure 3B).

**Figure 3:**
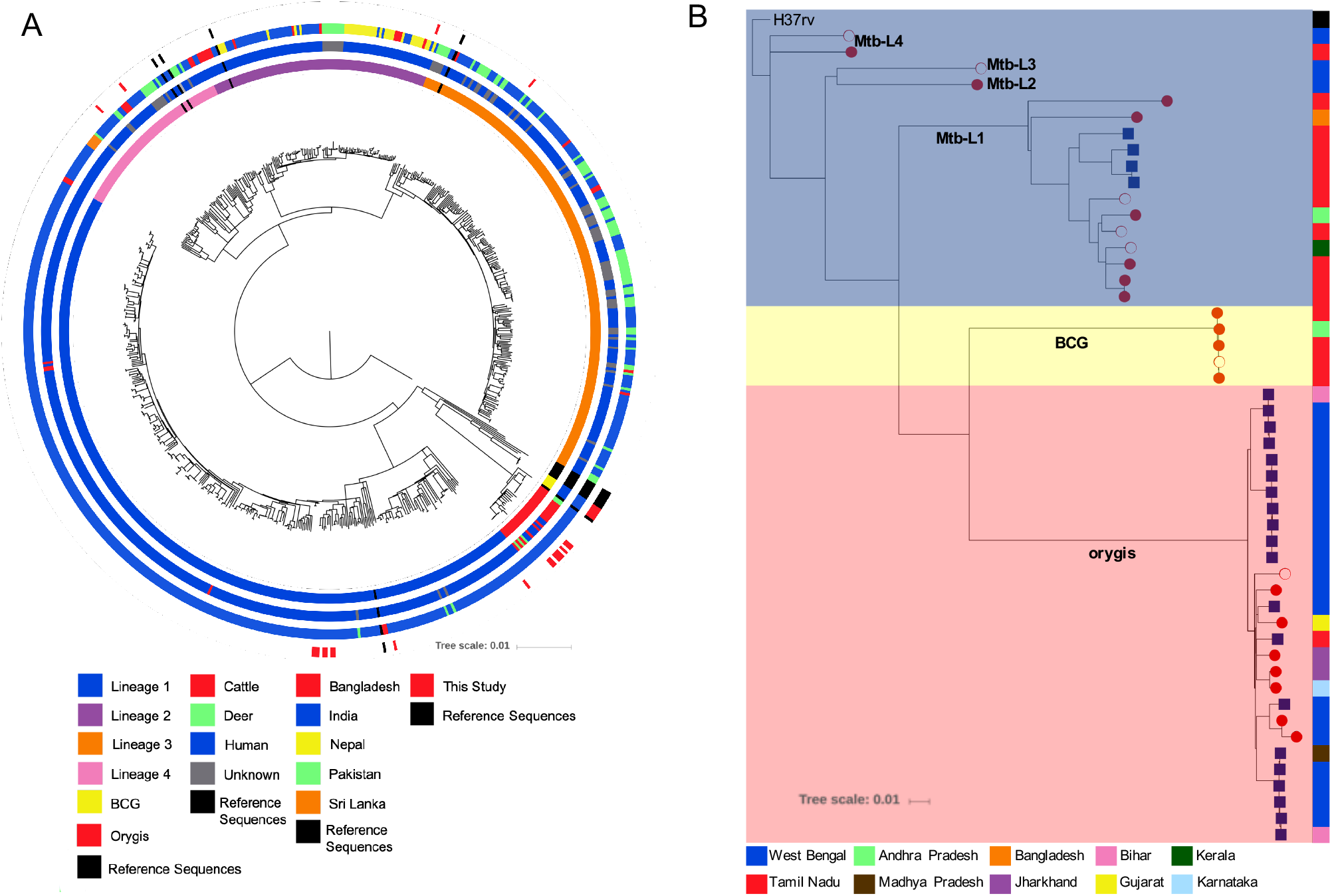
Phylogenies of newly sequenced isolates in the context of the MTBC isolates in South Asia Fig 3A: The circular tree shows the same samples (highlighted by red marks in the outer colored band) in the context of South Asia. The newly sequenced isolates were compared with 715 downloaded MTBC sequences downloaded from the SRA database. The sequences represent different lineages (inner colored bar): Mtb-L1 (blue), Mtb-L2 (purple), Mtb-L3 (orange), Mtb-L4 (pink), *M. bovis* BCG (yellow), *M. orygis* (red); different host species (second colored band from the tree): cattle (red), deer (green), human (blue), unknown (gray); different countries (third colored band from the tree): Bangladesh (red), India (blue), Nepal (yellow), Pakistan (green), Sri Lanka (orange). Note that cattle isolates included one bison isolate as a part of the Bovidae family. All MTBC lineage and species reference samples are shown in black throughout all variables in the metadata. Fig 3B: The isolates from this study, all downloaded *M. orygis* sequences and cattle lineage 1 sequences were included in the dendrogram. The background color represents the sequences identified as *M. tb* (blue), as *M. bovis* BCG (yellow), and as *M. orygis* (red). At the end of each node, the blue squares represent animal samples and the red circles represent human samples. The filled circles are extrapulmonary samples and the open circles are from pulmonary samples. The colored bar represents the states in India from where the sequences were collected: West Bengal (blue), Tamil Nadu (red), Andhra Pradesh (green), Madhya Pradesh (brown), Jharkhand (purple), Bihar (yellow), Gujarat (pink), Kerala (dark green) and Karnataka (light blue). One sample was from Bangladesh (orange).

## Discussion

Zoonotic TB is increasingly recognized as a potential threat to TB control (^23^). *M. bovis* was identified as a cause for bovine tuberculosis nearly a century prior to the description of other distinct MTBC subspecies infecting cattle, at which point the scientific community had arrived at a general consensus that zoonotic risks associated with TB were caused by *M. bovis* (^24^). This continues to be the definition of zTB provided by the WHO, OIE, and FAO today (^1,3,4^), despite mounting evidence that human infections can also be caused by other members of MTBC, such as *M. orygis* (^11–14^). In order to evaluate zTB prevalence in India, a surveillance screen of 940 clinical TB isolates from a large referral hospital in Southern India was performed. This search was then broadened to include 715 additional MTBC sequences deposited from South Asia available on the SRA database. Strikingly, no *M. bovis* was identified in this study. Several cases of TB were identified to be caused by *M. orygis*, and these were more likely to present as extrapulmonary TB. Together, these findings suggest that *M. bovis* may be an inadequate proxy for detecting zTB infection in countries such as India, and that *M. bovis* prevalence will underestimate the burden of zTB, especially in regions where *M. bovis* may not be the primary member or the MTBC circulating amongst livestock.

The detection of *M. orygis* in this study is concordant with previous reports linking *M. orygis* to patients from South Asia (^11–14^). Marcos *et al* identified 8 *M. orygis* isolates from humans originating from India, Pakistan, or Nepal (^12^). All 7 *M. orygis* isolates collected by Lavender *et al* were from patients from India (^13^). Moreover, *M. orygis* has also been detected in cattle and rhesus monkeys in Bangladesh (^11,14^). Collectively these data indicate that members of the MTBC complex, other than *M. bovis*, may be relatively more prevalent in the livestock in these geographies. Recently, Brites *et al* proposed that the distribution of *M. orygis* and *M. bovis* may be traced back to two independent cattle domestication events wherein *M. orygis* may have become a pathogen of cattle in South Asia, primarily *Bos indicus*, whereas *M. bovis* became a pathogen of *Bos taurus* (^25^). Virulence studies comparing pathogenicity between these two subspecies in infected cattle breeds have not yet been conducted. Alternatively, Rahim *et al* suggested that the global distribution of these subspecies indicates the emergence of *M. orygis* prior to *M. bovis* from a common MTBC ancestor, wherein *M. orygis* may have been dispersed to South Asia with the migration of humans and become established before the arrival of *M. bovis* (^14^). However, prior establishment of an MTBC lineage does not negate the introduction of another at a later point in time, as is demonstrated by the wide dispersal of *M. tuberculosis* lineages 2 and 4 (^26^).

*M. tuberculosis sensu stricto* isolates from South Asia were dispersed across lineages 1-4 with an enrichment of lineages 1 and 3, which is concordant with previous reports (^26^). Cattle origin MTBC sequences deposited from South Asia were distributed among *M. tuberculosis* lineage 1 and *M. orygis* lineages, highlighting the need to understand pathogenicity and transmission dynamics of these pathogens in cattle. There have been several reported cases of transmission of *M. tuberculosis* from humans into cattle, which may have important implications for transmission control in high TB endemic countries (^27,28^). Collectively, our data support the occurrence of zTB in India but suggest that this is associated with *M. orygis* (and possibly *M. tuberculosis*), rather than *M. bovis.* Further, WGS analysis indicated that RD12 may be an inadequate marker for detection of *M. tuberculosis*. This region was deleted in 2 *M. tuberculosis* isolates, and 7 *M. tuberculosis* isolates were categorized as inconclusive due to a SNP in the reverse primer sequence.

This study performed a screen of nearly one-thousand clinical TB isolates from positive MGIT® cultures. This provides a number of advantages compared to previously reported studies (^8–10^). The identification of isolates directly from broth cultures allows for the detection of active TB cases rather than indirect measures such as serology. The two-step protocol enabled provisional identification by PCR and assessment of inconclusive results by WGS. In addition, compared to conventional PCR, the five-probe multiplex real-time PCR allowed for streamlined differentiation of many MTBC subspecies in a single reaction (Supplementary Table 4), showing promise for easy adoption in routine zTB screening based on the needs and resources of the location. Furthermore, the detection of *M. bovis* BCG served as a positive control for finding *M. bovis*, were it present. However, the deletion of RD1, along with WGS analysis, confirmed that these isolates were BCG Russia, consistent with the currently used vaccine in India. An important limitation is that this is a single center surveillance study, therefore the isolates collected may be biased to locations within or nearby Tamil Nadu or to the patients who traveled to the study site (Figure 1). Future studies need to be conducted in other areas of India and other countries in South Asia in order to obtain a representative dataset.

In conclusion, this study indicates that *M. bovis* may be uncommon in India and therefore its detection is likely an ineffective proxy for zTB prevalence. The operational definition of zTB should be broadened to include other MTBC subspecies capable of causing human disease. The growing evidence supporting *M. orygis* endemicity in South Asia alongside the identification of *M. tuberculosis* in cattle highlights the importance of a One Health approach to TB control in India.

## Acknowledgements

This research was supported by the Bill & Melinda Gates Foundation (grant OPP1176950 to V.K.) and the Canadian Institutes for Health Research (grant FDN148362 to M.A.B). We would like to thank Shalini Elangovan and Deepa Mani for assistance in preparation of clinical samples. We would also like to thank Fiona McIntosh for technical support with assay development.

## References

1 Global tuberculosis report 2019. Geneva: World Health Organization; 2019. Licence: CCBY-NC-SA3.0IGO.

2 Waters WR, Palmer MV, Buddle BM, et al. Bovine tuberculosis vaccine research: historical perspectives and recent advances. Vaccine 2012; 30: 2611–22.

3 World Health Organization. Zoonotic Tuberculosis. 2017. Available online at: https://www.who.int/tb/areas-of-work/zoonotic-tb/ZoonoticTBfactsheet2017.pdf

4 World Health Organization. Roadmap for zoonotic tuberculosis. 2017. Available online at: https://www.oie.int/fileadmin/Home/eng/Our_scientific_expertise/docs/pdf/Tuberculosis/Roadmap_zoonotic_TB.pdf

5 World Health Organization. The End TB Strategy. 2014. Available online at: http://www.who.int/tb/strategy/End_TB_Strategy.pdf?ua=1.

6 United States Department of Agriculture. Livestock and Poultry: World Markets and Trade. 2019. Available online at: https://apps.fas.usda.gov/psdonline/circulars/livestock_poultry.pdf.

7 Srinivasan S, Easterling L, Rimal B, et al. Prevalence of Bovine Tuberculosis in India: A systematic review and meta-analysis. Transbound Emerg Dis 2018; 65: 1627–40.

8 Bapat PR, Dodkey RS, Shekhawat SD, et al. Journal of Epidemiology and Global Health Prevalence of zoonotic tuberculosis and associated risk factors in Central Indian populations. J Epidemiol Glob Health 2017; 7: 277–83.

9 Prasad HK, Singhal A, Mishra A, et al. Bovine tuberculosis in India: Potential basis for zoonosis. Tuberculosis 2005; 85: 421–8.

10 Shah NP, Singhal A, Jain A, et al. Occurrence of overlooked zoonotic tuberculosis: Detection of Mycobacterium bovis in human cerebrospinal fluid. J Clin Microbiol 2006; 44: 1352–8.

11 van Ingen J, Brosch R, van Soolingen D. Characterization of Mycobacterium orygis. Emerg Infect Dis 2013; 19: 521–2.

12 Marcos LA, Spitzer ED, Mahapatra R, et al. Mycobacterium orygis lymphadenitis in New York, USA. Emerg Infect Dis 2017; 23: 1749–51.

13 Lavender CJ, Globan M, Kelly H, et al. Epidemiology and control of tuberculosis in Victoria, a low-burden state in south-eastern Australia, 2005-2010. Int J Tuberc Lung Dis 2013; 17: 752–8.

14 Rahim Z, Thapa J, Fukushima Y, et al. Tuberculosis Caused by Mycobacterium orygis in Dairy Cattle and Captured Monkeys in Bangladesh : a New Scenario of Tuberculosis in South Asia. Transbound Emerg Dis 2017; 64: 1965–9.

15 Untergasser A, Cutcutache I, Koressar T, et al. Primer3 - new capabilities and interfaces. Nucleic Acids Res 2012; 40: e115.

16 Halse TA, Escuyer VE, Musser KA. Evaluation of a single-tube multiplex real-time PCR for differentiation of members of the Mycobacterium tuberculosis complex in clinical specimens. J Clin Microbiol 2011; 49: 2562–7.

17 Saïd-Salim B, Mostowy S, Kristof AS, et al. Mutations in Mycobacterium tuberculosis Rv0444c, the gene encoding anti-SigK, explain high level expression of MPB70 and MPB83 in Mycobacterium bovis. Mol Microbiol 2006; 62: 1251–63.

18 van Soolingen D, Hermans PW, de Haas PE, et al. Occurrence and stability of insertion sequences in Mycobacterium tuberculosis complex strains: Evaluation of an insertion sequence-dependent DNA polymorphism as a tool in the epidemiology of tuberculosis. J Clin Microbiol 1991; 29: 2578–86.

19 Kapur V, Li LL, Hamrick M, et al. Rapid Mycobacterium species assignment and unambiguous identification of mutations associated with antimicrobial resistance in Mycobacterium tuberculosis by automated DNA sequencing. Arch Pathol Lab Med 1995; 119: 131–8.

20 Stamatakis A. RAxML version 8: A tool for phylogenetic analysis and post-analysis of large phylogenies. Bioinformatics 2014; 30: 1312–3.

21 Gardner SN, Slezak T, Hall, BG. kSNP3.0: SNP detection and phylogenetic analysis of genomes without genome alignment or reference genomes. Bioinformatics 2015; 31: 2877–8.

22 Letunic I, Bork P. Interactive Tree Of Life (iTOL) v4: recent updates and new developments. Nucleic Acid Res 2019; 47:W256–9.

23 Olea-Popelka F, Muwonge A, Perera A, et al. Zoonotic tuberculosis in human beings caused by Mycobacterium bovis — a call for action. Lancet Infect Dis 2017; 17: e21–5.

24 Smith T. A comparative study of bovine tubercle bacilli and of human bacilli from sputum. J Exp Med 1898; 3: 451–511.

25 Brites D, Loiseau C, Menardo F, et al. A new phylogenetic framework for the animal-adapted Mycobacterium tuberculosis complex. Front Microbiol 2018; doi:10.3389/fmicb.2018.02820.

26 Wiens KE, Woyczynski LP, Ledesma JR, et al. Global variation in bacterial strains that cause tuberculosis disease: a systematic review and meta-analysis. BMC Med 2018; 16: 196.

27 Sweetline Anne N, Ronald BS, Kumar TM, et al. Molecular identification of Mycobacterium tuberculosis in cattle. Vet Microbiol 2017; 198: 81–7.

28 Ocepek, M, Pate M, Zolnir-Dovc M et al. Transmission of Mycobacterium tuberculosis from Human to Cattle. J Clin Microbiol 2005; 43: 3555–7.

