## Supplementary Material for "Zoonotic tuberculosis in India: looking beyond *Mycobacterium bovis*"

Supplementary data

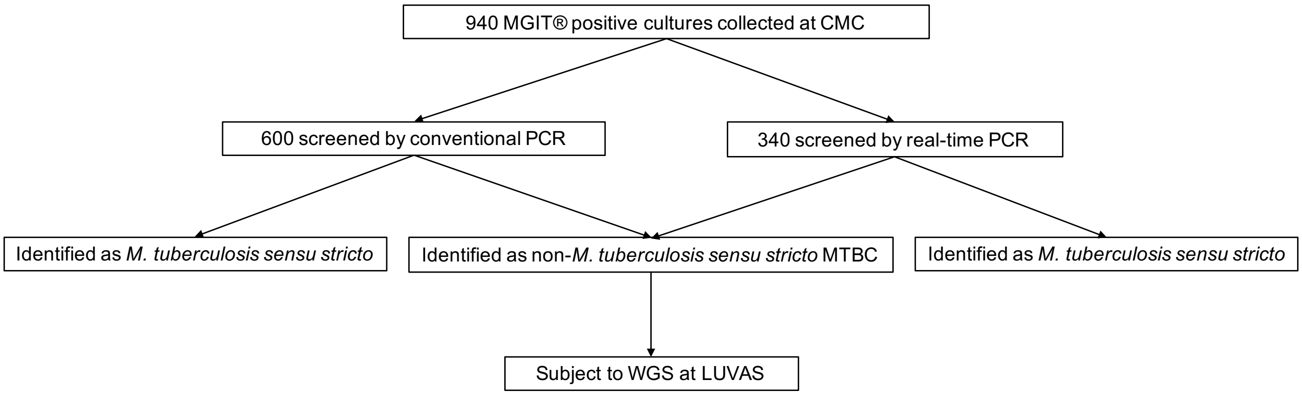


***Supplementary Figure 1:*** **Study design**

| **Sample type** | **Number** |
|---|---|
| Abdomen/Peritoneal fluid | 15 |
| Anal fistula | 2 |
| Biopsy | 5 |
| Bone | 43 |
| Bone marrow | 2 |
| Brain abscess | 1 |
| Colon | 5 |
| Cerebrospinal fluid | 27 |
| Fluid | 5 |
| Lymph node | 162 |
| Muscle abscess | 5 |
| Pus | 56 |
| Pericardium | 1 |
| Skin | 1 |
| Synovium | 8 |
| Tissue | 40 |
| Urine | 13 |
| Unspecified | 1 |
| Total | 392 |

***Supplementary Table 1:*** **Number and tissue types of extrapulmonary samples**
Extrapulmonary samples were defined as cultures from tissues other than the lungs or lung fluid.

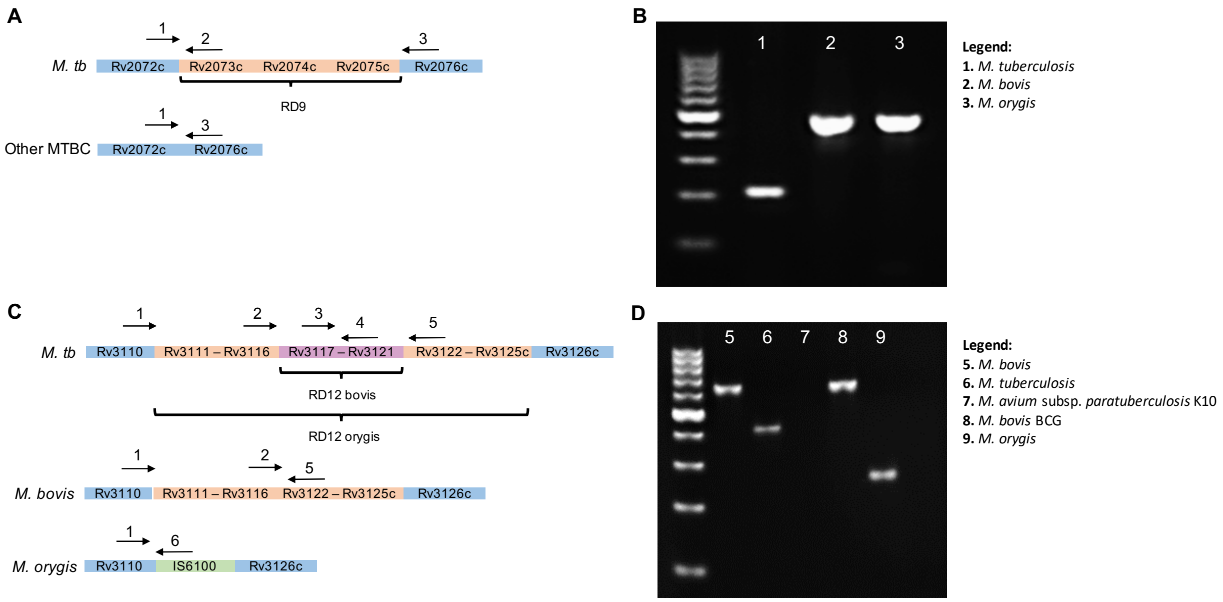


***Supplementary Figure 2:*** **Conventional PCR to detect differences in deletions of RD9 and RD12**

**SFig 2A:** A three-primer PCR reaction was developed to detect the presence of absence of RD9, which is found in *M. tb* but is absent in all other MTBC members. **SFig 2B:** A 209bp band is amplified when *M. tb* is present and a 410bp band is amplified when other MTBC members are present. **SFig 2C:** A six-primer PCR was developed to detect differences in the deletion size of RD12. RD12 is present in *M. tb* and absent in *M. bovis* and *M. orygis.* In *M. orygis,* the RD12 deletion is larger and is replaced with IS6100. **SFig 2D:** A 409bp band is amplified when *M. tb* is present, a 615bp band is amplified when *M. bovis* or BCG is present, and a 264bp band is amplified when *M. orygis* is present.

| Assay | Name | Sequence (5' → 3') | Dye | Quencher |
|---|---|---|---|---|
| Conventional PCR | RD9_Forward | CCGATACCATGCAACAACGG | | |
| | RD9_Reverse1 | CGGTCTCTCCGAGCATTC | | |
| | RD9_Reverse2 | GCTCGAGCTAGACCTGCAC | | |
| | RD12Mtb_Forward | GTATTTGCGCCCATATCCTGG | | |
| | RD12Mtb_Reverse | CCTGGCTTCAAGCACCATTC | | |
| | RD12Mbovis_Forward | GGCCATCAACGTCAAGAACCTC | | |
| | RD12Mbovis_Reverse | CGAACTCGTATTTTGTGGCCAC | | |
| | RD12Morygis_Forward | GTGGAAATGGAAGCGTTGACC | | |
| | RD12Morygis_Reverse | GGTACCTCCTCGATGAACCAC | | |
| Real-time PCR | Rv0444c_Probe | CTCGGCTGACCCGA | FAM | MGBNFQ |
| | Rv0444c_Forward | GATGCTGGGCACCATTGTC | | |
| | Rv0444c_Reverse | GCCCACCGGTACCATCTTG | | |
| | RD1 Probe | CACTCTGAGAGGTTGTCA | VIC | MGBNFQ |
| | RD1_Forward | CCCTTTCTCGTGTTTATACGTTTGA | | |
| | RD1_Reverse | GCCATATCGTCCGGAGCTT | | |
| | RD9 Probe | AGGTTTCA+CCTTCGAC+CC | TEX615 | BHQ |
| | RD9_Forward | TGCGGGCGGACAACTC | | |
| | RD9_Reverse | CACTGCGGTCGGCATTG | | |
| | RD12 Probe | TGCGCTGACCCCAC | NED | MGBNFQ |
| | RD12_Forward | CGTTGGAACGCGAAATACG | | |
| | RD12_Reverse | CCAGGATATGGGCGCAAAT | | |
| | extRD9_Probe | G+TT+CTTCAG+CTGGT+CC | CY5 | BHQ |
| | extRD9_Forward | GCCACCACCGACTCATAC | | |
| | extRD9_Reverse | CGAGGAGGTCATCCTGCTCTA | | |

***Supplementary Table 2:*** **Primer and probe sequences for conventional and real-time PCR assays**

The real-time PCR RD1, RD9, RD12, and ext-RD9 probes and primers are as described in Halse *et al*. The Rv0444c, RD1, and RD12 probes are Taqman MGB probes. The RD9 and ext-RD9 probes are locked nucleic acid probes. A '+' indicates insertion of a locked nucleic acid base.

| Assay | Master mix | Thermocycling conditions |
|---|---|---|
| Conventional PCR | 6.25 µl of 10X *Taq* buffer (Thermo Scientific)<br>6.25 µl acetamide 50% (wt/vol)<br>1.6 mM $MgCl_2$<br>0.2 mM deoxynucleoside triphosphates (dNTPs)<br>2.5 U per reaction *Taq* polymerase (Thermo Scientific)<br>500 nM of each primer<br>5µl of template DNA<br>Sterile water<br>50µl total volume | Initial denaturation 94°C for 3 minutes<br>35 cycles of:<br>- Denaturation at 94°C for 30 seconds<br>- Annealing at 55°C for 1 minute<br>- Elongation at 72°C for 1 minute,<br>Final elongation step at 72°C for 10 minutes |
| Real-time PCR | 10 µl TaqMan multiplex master mix (Applied Biosystems)<br>450 nM of each primer<br>125 nM of each probe<br>1 µl of template DNA<br>Sterile water<br>20µl total volume | 95°C for 10 minutes<br>40 cycles of:<br>- 95°C for 15s<br>- 60°C for 1 minute |

***Supplementary Table 3:*** **Master mix preparation and thermocycling conditions for conventional and real-time PCR assays**

| | RD1 | RD9 | RD12 | Rv0444c | Ext-RD9 |
|---|---|---|---|---|---|
| ***M. tuberculosis*** | + | + | + | - | + |
| ***M. orygis*** | + | - | - | + | + |
| ***M. bovis*** | + | - | - | - | + |
| ***M. bovis* BCG** | - | - | - | - | + |
| ***M. africanum*** | + | - | + | - | + |
| ***M. microti*** | - | - | + | - | + |
| **NTMs** | - | - | - | - | - |

***Supplementary Table 4:*** **Interpretation of RT-PCR results to determine MTBC sample identity**

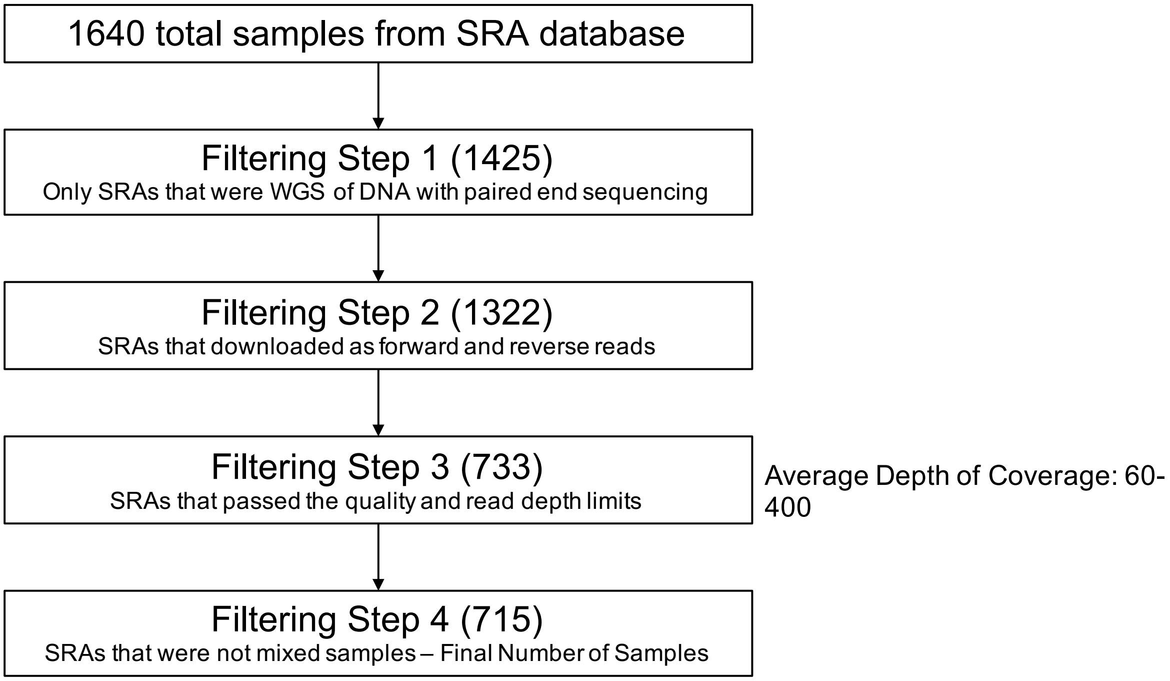


***Supplementary Figure 3:*** **Selection and filtering pipeline of downloaded SRA MTBC genomes from South Asia**
A total of 1640 SRAs were downloaded from NCBI with the search terms ("*Mycobacterium bovis*" OR "*Mycobacterium tuberculosis*" OR "*Mycobacterium africanum*" OR "*Mycobacterium orygis*" OR "*Mycobacterium canetti*" OR "*Mycobacterium caprae*" OR "*Mycobacterium bovis* BCG" NOT "H37Rv" NOT "H37Ra") from ("India" OR "Bangladesh" OR "Nepal" OR "Sri Lanka" OR "Pakistan"). Through multiple filtering steps, the total number of SRAs analyzed in the phylogenetic tree in Fig 2B was 715. The lists of SRAs after each filtering step can be found in the Supplemental Tables 4-8.

***Supplementary Tables 5 – 7*****: Uploaded as Excel files**

| Isolate number | Identification by real-time PCR | Average coverage | Genome coverage (%) | Phred quality score R1 | Phred quality score R2 | SNP count | Identification by WGS |
|---|---|---|---|---|---|---|---|
| P70 | *M. bovis* BCG | 103.2 | 99.35 | 34.6 | 31.1 | 803 | *M. bovis* BCG Russia |
| P280 | *M. bovis* BCG | 80.4 | 99.15 | 34.5 | 31.1 | 1834 | *M. tuberculosis* lineage 2 |
| E50 | *M. bovis* BCG | 54.4 | 99.49 | 33.7 | 30.3 | 784 | *M. bovis* BCG Russia |
| E110 | *M. bovis* BCG | 85.0 | 99.68 | 33.8 | 29.6 | 797 | *M. bovis* BCG Russia |
| E280 | *M. bovis* BCG | 77.2 | 99.33 | 34.1 | 28.0 | 804 | *M. bovis* BCG Russia |
| E396 | *M. bovis* BCG | 98.7 | 99.38 | 33.9 | 28.8 | 817 | *M. bovis* BCG Russia |
| P326 | Inconclusive | 53.5 | 99.27 | 33.9 | 29.2 | 2322 | *M. tuberculosis* lineage 1 |
| P414 | Inconclusive | 82.2 | 99.42 | 33.6 | 27.8 | 2412 | *M. tuberculosis* lineage 1 |
| P448 | Inconclusive | 78.7 | 99.36 | 34.8 | 30.3 | 987 | *M. tuberculosis* lineage 4 |
| P465 | Inconclusive | 121.2 | 99.46 | 34.1 | 28.7 | 2476 | *M. tuberculosis* lineage 1 |
| E343 | Inconclusive | 277.6 | 99.53 | 35.1 | 32.5 | 2607 | *M. tuberculosis* lineage 1 |
| E369 | Inconclusive | 75.1 | 99.33 | 33.8 | 29.5 | 2429 | *M. tuberculosis* lineage 1 |
| E379 | Inconclusive | 115.2 | 99.51 | 34.2 | 29.3 | 2499 | *M. tuberculosis* lineage 1 |
| E415 | Inconclusive | 153.8 | 99.53 | 33.9 | 28.9 | 2510 | *M. tuberculosis* lineage 1 |
| E277 | *M. tuberculosis* | 138.6 | 99.39 | 34.8 | 30.4 | 1691 | *M. tuberculosis* lineage 3 |
| E428 | *M. tuberculosis* | 105.6 | 99.66 | 34.0 | 29.0 | 2594 | *M. tuberculosis* lineage 1 |
| P429 | *M. orygis* | 12.2 | 97.49 | 34.3 | 30.7 | 2299 | *M. orygis* |
| E36 | *M. orygis* | 72.9 | 98.26 | 33.5 | 28.2 | 2547 | *M. orygis* |
| E65 | *M. orygis* | 34.6 | 97.75 | 33.4 | 29.2 | 2408 | *M. orygis* |
| E120 | *M. orygis* | 11.1 | 97.15 | 33.9 | 31.1 | 2330 | *M. orygis* |
| E157 | *M. orygis* | 64.7 | 98.26 | 34.1 | 28.9 | 2478 | *M. orygis* |
| E313 | *M. orygis* | 58.8 | 98.28 | 34.1 | 31.6 | 2483 | *M. orygis* |
| E374 | *M. orygis* | 139.1 | 98.45 | 34.6 | 31.1 | 2566 | *M. orygis* |
| E186 | *M. tuberculosis* RD12 absent | 90.3 | 98.74 | 34.1 | 29.1 | 968 | *M. tuberculosis* lineage 4 |
| E363 | *M. tuberculosis* RD12 absent | 107.4 | 99.00 | 34.5 | 30.2 | 2474 | *M. tuberculosis* lineage 1 |

***Supplementary Table 8:*** **Selection, library preparation and whole genome sequencing data of 25 selected isolates**

| Sample name | PCR ID | Top BLAST match | Seq length | Query coverage | % Identity |
|---|---|---|---|---|---|
| E133 | NTM | *Mycobacterium phocaicum* | 430 | 100.00% | 99.77% |
| E153 | NTM | *Mycobacterium engbaekii* | 440 | 99.00% | 99.32% |
| E193 | NTM | *Mycobacterium abscessus* | 440 | 100.00% | 100.00% |
| P24 | NTM | *Mycobacterium sp. K328YA* | 438 | 94.00% | 100.00% |
| P30 | NTM | *Mycobacterium alvei* | 437 | 100.00% | 98.40% |
| P390 | NTM | *Mycobacterium abscessus* | 432 | 100.00% | 100.00% |
| P427 | NTM | *Mycobacterium intracellulare* | 419 | 100.00% | 100.00% |
| P146 | NTM | *Mycobacterium simiae* | 443 | 99.00% | 99.32% |
| P149 | NTM | *Mycobacterium intracellulare* | 424 | 100.00% | 99.76% |
| E22 | NTM | *Mycobacterium abscessus* | 357 | 100.00% | 99.72% |
| P219 | NTM | *Mycobacterium yongonense* | 359 | 100.00% | 99.44% |
| P281 | NTM | *Mycobacterium fortuitum* | 362 | 100.00% | 100.00% |
| P426 | NTM | *Mycobacterium intracellulare* | 333 | 100.00% | 100.00% |
| P81 | NTM | *Mycobacterium parascrofulaceum* | 381 | 100.00% | 99.48% |

***Supplementary Table 9:*** **Hsp65 sanger sequencing results of non-tuberculous mycobacteria (NTM) isolates**

| | P429 | E120 | E36 | E374 | E313 | E65 | E157 |
|---|---|---|---|---|---|---|---|
| P429 | 0 | 85.47 | 88.27 | 89.10 | 89.66 | 88.84 | 89.52 |
| E120 | 282 | 0 | 86.30 | 87.55 | 88.13 | 87.27 | 87.84 |
| E36 | 271 | 285 | 0 | 91.53 | 92.02 | 91.19 | 91.77 |
| E374 | 252 | 259 | 214 | 0 | 93.29 | 92.38 | 93.07 |
| E313 | 239 | 247 | 202 | 170 | 0 | 92.93 | 93.64 |
| E65 | 258 | 265 | 223 | 193 | 177 | 0 | 97.39 |
| E157 | 242 | 253 | 208 | 175 | 161 | 66 | 0 |

***Supplementary Figure 4*: SNP distances between 7 *M. orygis* isolates from this study**
The intensity of the color corresponds to the distance between isolates. The bottom portion of the matrix indicates the number of SNPs between the two isolates. The top portion indicates the percent of total SNPs shared between them

***Supplementary Table 10*: Uploaded as Excel file**

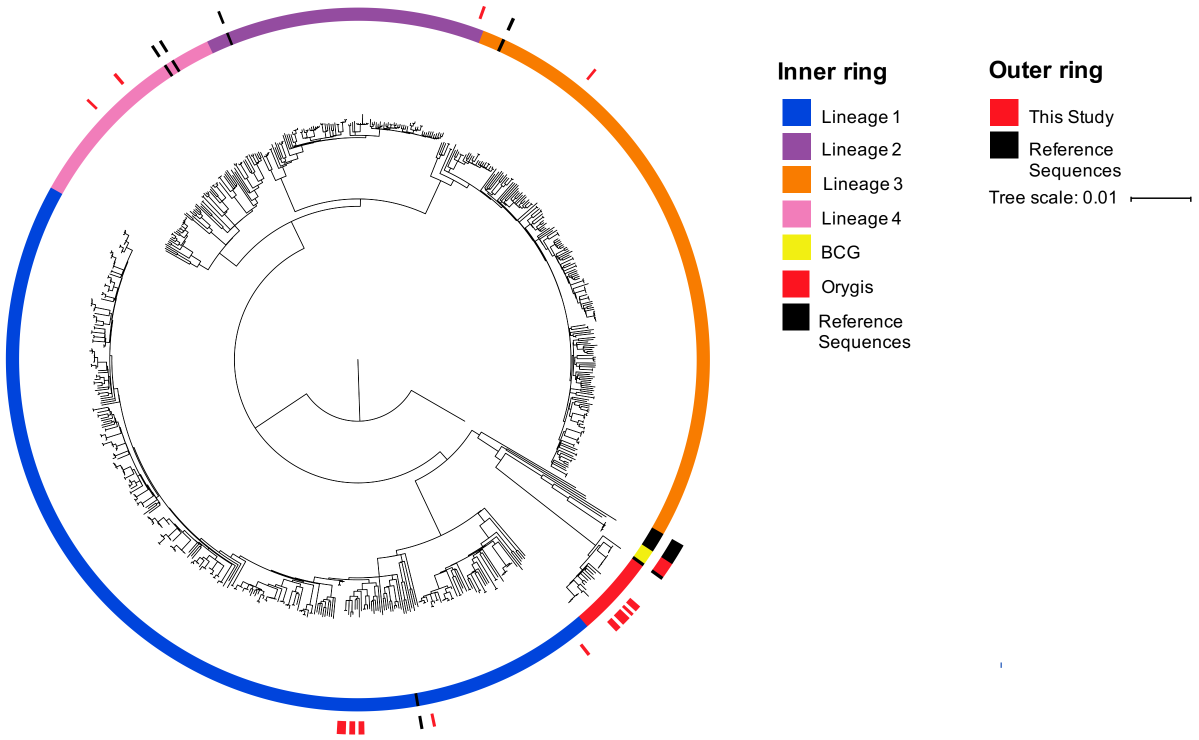


***Supplementary Figure 5:*** **Maximum likelihood phylogenetic tree of newly sequenced isolates and 715 MTBC genomes collected from South Asia with lineage metadata**

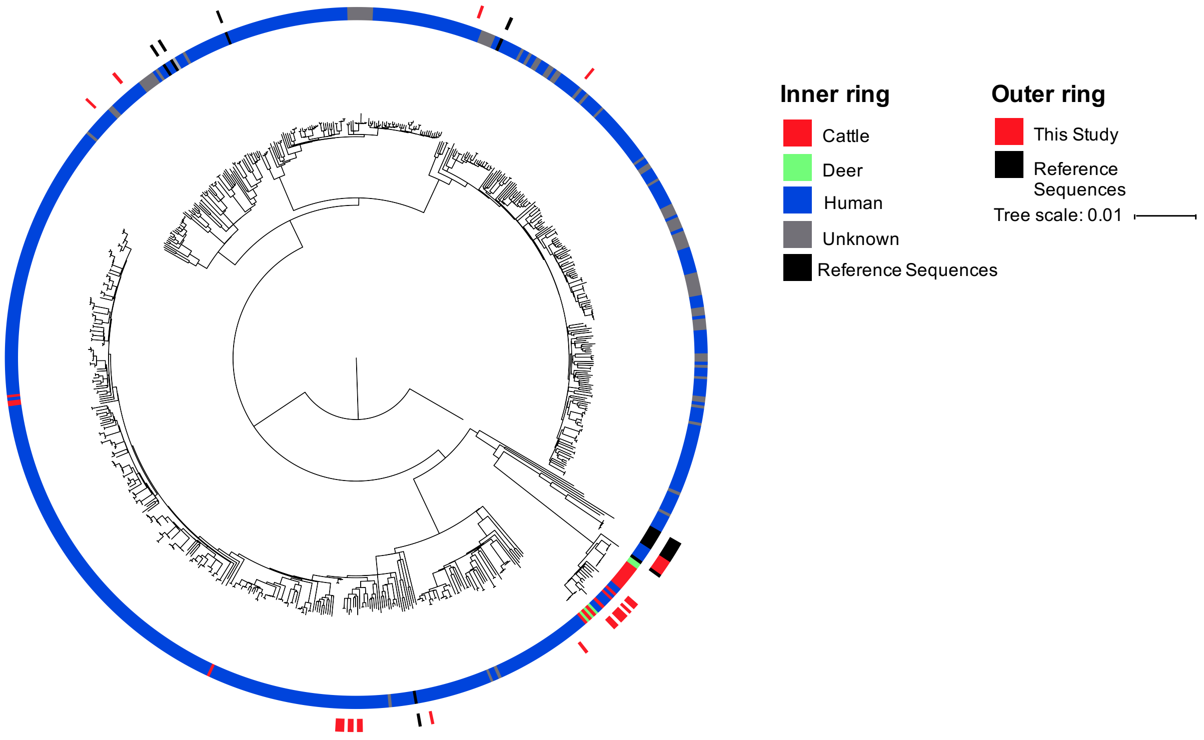


***Supplementary Figure 6:*** **Maximum likelihood phylogenetic tree of newly sequenced isolates and 715 MTBC genomes collected from South Asia with host metadata**

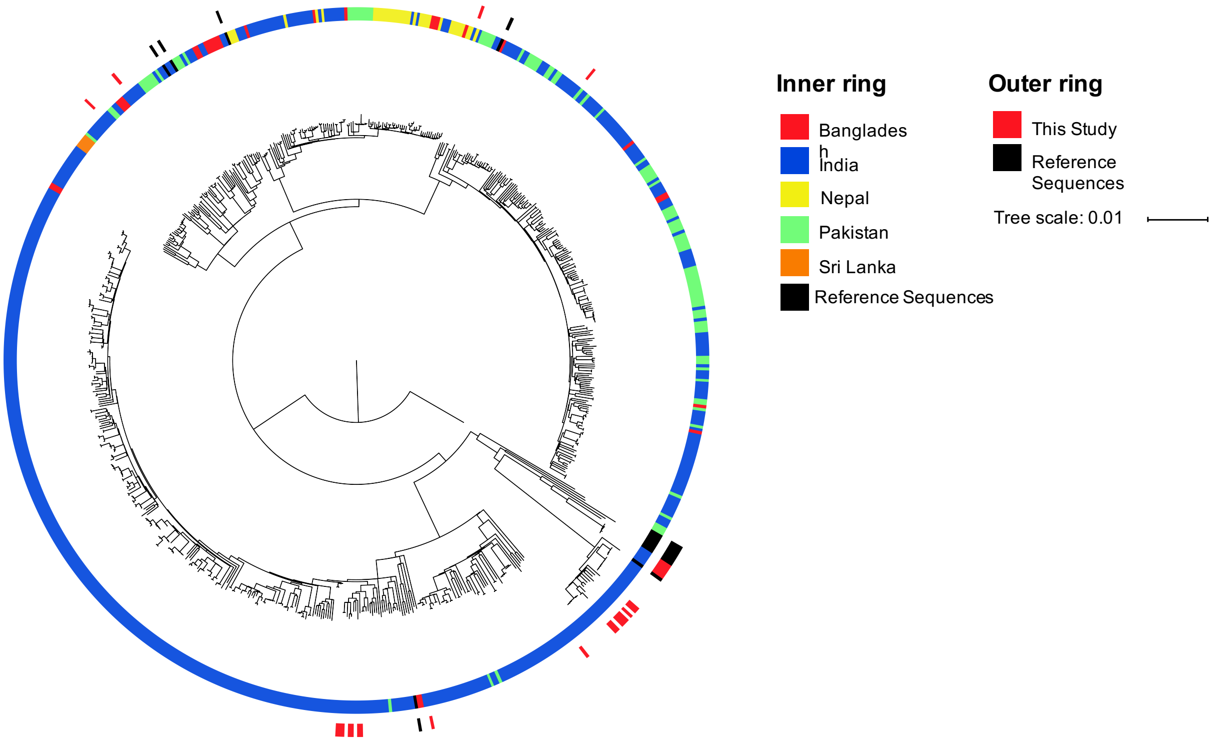


***Supplementary Figure 7:*** **Maximum likelihood phylogenetic tree of newly sequenced isolates and 715 MTBC genomes collected from South Asia with country metadata**

**Supplementary Methods**

*Bioinformatics*

Sequences were assessed using the United States Department of Agriculture Animal and Plant Health Inspection Service Veterinary Services pipeline vSNP (https://github.com/USDA-VS/vSNP). The vSNP pipeline involved a two-step process. Step 1 determined SNP positions called within the sequence. Paired FASTQ files were processed using BWA-MEM to align reads to a reference genome *M. tuberculosis* H37Rv (NC_000962.3) for sequences included in this study ([1]). Duplicate reads were tagged and removed using the Mark Duplicates tool from Picard v 2.20.2 (http://broadinstitute.github.io/picard). SNPs were called using FreeBayes v. 1.3.1 ([2]). Unmapped reads shorter than 64 base pairs were removed and low-coverage contigs with an average k-mer coverage of less than 5 were removed. Depth of coverage was calculated using Pysam (https://github.com/pysam-developers/pysam) and positions with zero coverage were added to the VCF file. Step 2 assessed SNPs called between closely related isolate groups to output SNP alignments, tables and phylogenetic trees. For a SNP to be considered in a group there must have been at least one position with an allele count (AC) =2, quality score >150 and map quality > 56. Once determined, SNPs were aligned using the following workflow. If the quality score for a SNP position was greater than 150, the alternate allele was called if AC=2. However, if AC=1, the position was called ambiguous. Deletions were called when the alternate allele was a gap. If the quality score was between 50 and 150, the allele was marked N. If the quality score was less than 50, then the reference allele was called. Uninformative SNPs were not included. BAM files were used to visualize SNP calls. Unreliable positions due to read alignment error may have been removed from the analysis. The output SNP alignment was used to assemble a maximum likelihood phylogenetic tree using RAxML ([3]).

*Phylogenetic tree assembly*

To compare the 25 newly sequenced genomes in the context of sequences from South Asia, a NCBI Sequence Read Archive (SRA) (https://www.ncbi.nlm.nih.gov/sra) search was performed using the search terms ("*Mycobacterium bovis*" OR "*Mycobacterium tuberculosis*" OR "*Mycobacterium africanum*" OR "*Mycobacterium orygis*" OR "*Mycobacterium canetti*" OR

"*Mycobacterium caprae*" OR "*Mycobacterium bovis* BCG" NOT "H37Rv" NOT "H37Ra") from ("India" OR "Bangladesh" OR "Nepal" OR "Sri Lanka" OR "Pakistan"). This search yielded 1640 genomes. These sequences were then filtered prior to tree assembly (Supplementary Figure 3). A total of 215 were excluded because they were not from studies that performed with paired end sequencing. A further 103 were excluded as they did not contain forward and reverse read files. Another 589 sequences whose average depth of coverage was not between 60-400 were excluded. Finally, 18 were excluded after running vSNP step 2 due to the samples being mixed and generating multiple SNPs at all locations in the genome. In total, 715 sequences remained (Supplementary Table 5). All sequences were download from the SRA using the fasterq-dump tool from the sra toolkit v. 2.9.6 (https://ncbi.github.io/sra-tools/) and sequences were run through steps 1 and 2 of vSNP. Phylogenetic trees were constructed using vSNP to compare the 25 sequences from this study with the total 715 available genomes from South Asia and a subset. Reference sequences were also included for comparison (Supplementary Table 6). Phylogenetic trees were rooted to *M. tuberculosis* H37Rv. To compare the sequences collected in this study in the context of the global MTC, treeSPAdes (http://cab.spbu.ru/software/spades/) was used to assemble reads for kSNP3 ([4]). Genomes assemblies had expected complete genome sizes. The kSNP3 manual instructions were followed using kchooser calculated kmer value. All phylogenetic trees were visualized using the Interactive Tree of Life (iTOL) with their respective metadata (Supplementary Table 7) ([5]).

References:

1 Li H. Aligning sequence reads, clone sequences and assembly contigs with BWA-MEM. *arXiv e-prints* 2013; arXiv:1303.3997

2 Garrison E, Marth G. Haplotype-based variant detection from short-read sequencing. *arXiv e-prints* 2012; arXiv:1207.3907

3 Stamatakis A. RAxML version 8: A tool for phylogenetic analysis and post-analysis of large phylogenies. *Bioinformatics* 2014; **30**: 1312–3.

4 Gardner SN, Slezak T, Hall, BG. kSNP3.0: SNP detection and phylogenetic analysis of genomes without genome alignment or reference genomes. *Bioinformatics* 2015; **31**:

2877-8.

5 Letunic I, Bork P. Interactive Tree Of Life (iTOL) v4: recent updates and new developments. *Nucleic Acid Res* 2019; **47:** W256-9.
